# Internal Resonance Dynamics in a Delayed van der Pol Oscillator Modeling Basal Ganglia Oscillations

**DOI:** 10.1101/2025.06.16.660028

**Authors:** M. A. Elfouly, T. S. Amer

## Abstract

Movement disorders, like Parkinson’s disease, happen because of unusual patterns in the connections between the cortex and basal ganglia, often caused by timing issues in feedback pathways. This study uses a two-delay nonlinear dynamic model, based on the delayed the van der Pol oscillator, to examine how delays in the feedback loops of the direct and indirect basal ganglia pathways lead to unusual movement patterns and resonance. A thorough analysis of resonance looks at how the ratios of delays and the time it takes to respond affect the system’s frequency, showing when internal resonance occurs and how frequency stabilizes under different conditions. We examine how stable the system is by looking at changes in its behavior and measuring the Lyapunov exponent across distinct types of nonlinear feedback setups. Simulations demonstrate transitions from stable oscillations to chaos with varying delays and saturation strength. Our results reproduce symptoms of Parkinson’s disease, such as resting tremor, dyskinesia, and freezing, demonstrating how delayed inhibition or hyperactivity destabilizes motor function. The discussion supports these findings, indicating that early problems start in the striatum, with complex effects in the globus pallidus that worsen motor symptoms. This model explains the temporal evolution of Parkinson’s disease symptoms and highlights the timing of feedback and saturation as key therapeutic targets. Overall, this research offers a biological explanation for motor problems caused by delays and supports novel approaches for brain stimulation using flexible methods.

## 1. Introduction

Movement disorders, like Parkinson’s disease, happen because of problems with how neurons communicate and coordinate in the basal ganglia-cortical circuits. This study aims to explain how these timing problems, known as dual feedback delays, lead to harmful rhythms like resting tremor or bradykinesia. We employ a modified van der Pol (VdP) oscillator to elucidate these nonlinear dynamics and investigate their ramifications for deep brain stimulation therapy.

Dynamical systems are essential tools in natural sciences, engineering, and applied mathematics, offering a clear way to study how systems change over time due to both internal and external factors. These systems demonstrate a wide variety of behaviors, ranging from stable equilibrium points and periodic oscillations to quasi-periodic motions and dynamic chaos [1–4]. Nonlinear systems add extra complexity because their interactions change over time and are not always proportional, which can cause ongoing oscillations that don’t fade away on their own. Dynamic modeling is now essential in many fields: in physics, it helps explain how objects move when forces like friction and gravity act on them; in engineering, it assesses how stable mechanical and electronic systems are; and in neuroscience, it studies how nerve signals and movement control work together [5–9].

Time delay is an important factor that greatly affects how nonlinear systems behave by creating memory effects, which connect the current state of a system to its past behavior. Time delays naturally arise across a wide spectrum of natural and engineered systems, including neural networks, biological circuits, control systems, and communication infrastructures [10–12]. In systems that rely on feedback, delays can make the system unstable, causing sudden changes between stable states, starting oscillations, or even leading to chaotic behavior [13–15]. Mathematically, time delays change the frequency patterns and make it harder to analyze oscillations compared to systems that respond immediately. In engineering applications, such as adaptive control systems, poorly regulated delays can induce instability and performance degradation [16–18]. In biological systems, time delays are crucial for controlling how signals are sent in the brain and for creating the regular patterns needed for movement and thinking [19–21]. In brain circuits, having the right timing between neural signals is crucial for keeping activities in sync; if this timing is off, it can lead to unhealthy brain rhythms [22–29]. Therapies like deep brain stimulation (DBS) can help fix timing problems in the brain, which helps restore normal brain activity and reduce symptoms. A deeper understanding of how time delays affect brain rhythms gives researchers important information about the causes of neurological disorders and helps in creating more focused and effective treatments [30–32].

Resonance is a fundamental phenomenon in dynamic systems. It transpires when the system’s intrinsic frequencies synchronize with an internal or external stimulus, hence enhancing the oscillatory response, which can significantly influence the system’s stability and dynamic behavior. This phenomenon occurs when the system is excited at a frequency that coincides with one of its natural frequencies, amplifying vibrations and oscillations. This effect can make the system unstable or even fail [33–36]. Engineers have observed this in situations where wind or repetitive loads cause bridges to collapse due to resonant frequencies [37-38]. Resonance is an important phenomenon in biological and neural systems. It has a significant impact on brain oscillations and neural synchronization, which are associated with motor and cognitive functions [39-42]. It is also used in medicine, for example, in DBS for the treatment of neurological disorders [43].

Resonance is classified into two basic types: external resonance and internal resonance, both of which affect system dynamics in distinct ways. External resonance occurs when an external force excites a system of a similar frequency or approaches one of its intrinsic frequencies. This amplifies the oscillatory response. This phenomenon is used in various applications, including the creation of electrical filters, the investigation of mechanical vibrations, and the calibration of acoustic equipment [44].

On the other hand, internal resonance occurs when the system transfers energy between its inherent frequencies without external stimuli, resulting in the formation of new oscillation patterns. It is more complex, as it depends on nonlinear interactions between different internal modes of oscillation [45–47]. These interactions can result in complex transitions, such as shifts from periodic motion to chaos or desynchronization that alters system stability. In nonlinear systems, internal resonance is especially important for identifying oscillatory behavior, as it can either amplify or suppress oscillations depending on how internal frequency relationships are configured [48–50]. Its influence spans various scientific and technological domains, where it may lead to new oscillatory modes, internal energy redistribution, or sudden changes in dynamic behavior [51–54].

When time delays interact with neural feedback, they can generate internal oscillatory loops that maintain rhythmic activity even in the absence of external stimuli. The evidence suggests that internal resonance may contribute significantly to the persistence of pathological brain oscillations [55–58]. Specifically, small differences in the time it takes for signals to travel in feedback loops can greatly affect the natural frequencies of neural networks, either making oscillation patterns more stable or unstable [59–60]. Figuring out how time delays change the way brain rhythms work can lead to new ways to help fix healthy brain activity using treatments like electrical or magnetic stimulation [61–64]. We commonly use delay differential equations to mathematically describe how delayed feedback affects interactions and synchronization in brain circuits [65–69]. To systematically study the influence of delayed feedback on oscillatory behavior, we adopted a modified classic VdP oscillator, incorporating two distinct time delays. This model was chosen because it shows key nonlinear features found in unusual neural oscillations, like steady oscillations, limit cycle behavior, and how amplitude relates to frequency. Furthermore, its mathematical simplicity enables detailed exploration of time delay effects, which can be difficult with more complex biophysical neuronal models [70–72].

The VdP oscillator is a key model for examining how two delays affect the stability and frequency of brain networks that oscillate, particularly in conditions like Parkinson’s disease.

In its classical form, it is expressed as:

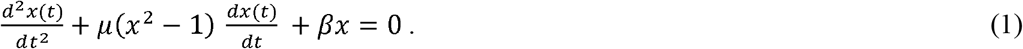

where *x*(*t*) is the state variable describing the system, *μ* is the nonlinearity parameter and *β* is the damping coefficient.

In the delayed form, the system is described by the following differential equation:

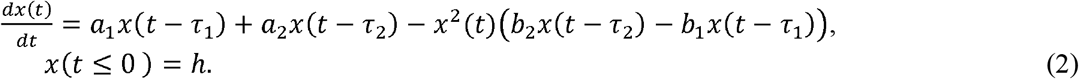

where *x*(*t*) state variable describing the system, *a*_1_ and *a*_2_ are the strengths of delayed feedback from two pathways, and for the delayed feedback from another pathway, *τ*;_1_ and *τ*;_2_ represent the response times of signal transmission in the first and second feedback loops and _*b*1_ and *b*_2_ commonly represent nonlinear interaction strengths between the current state *x*(*t*) and the delayed signals.

In this model, the parameters *a*_1_ and *a*_2_ show how strong the delayed feedback is from two key neural pathways in the basal ganglia: the direct pathway and the indirect pathway. The negative signals *a*_1_ and *a*_2_ represent the excitation and inhibition of motor cortex activity. In the mathematical model, the nonlinear coefficients *b*_1_ and *b*_2_ show how feedback changes based on the square of the activity level *x*^2^(*t*), meaning that the feedback effect gets stronger as activity increases. Negative values show that too much activity is being suppressed, a phenomenon referred to as “saturation” or “nonlinear inhibition”. It ascertains the impact of the nonlinear feedback signal on the system’s behavior. When *b*_1_ < 0, this term functions as an inhibitory force (negative saturation); when *b*_1_ > 0, it transforms into a facilitating mechanism (positive saturation). The same principle applies to *b*_2_ in the indirect pathway when *b*_2_ < 0; this term functions as an inhibitory force (negative saturation).

A specific set of values was chosen to illustrate the interaction of motor control; a specific set of values was selected to show how excitement, inhibition, and saturation interact in the cortical-basal ganglia loops. These states include resting, hyperactivity, inhibition, and a fixed time delay (*τ*_1_). Delay *τ*_1_ is kept small to capture rapid cortical feedback, with *τ*_2_ as a multiple to simulate the delays associated with the indirect pathway.

At rest, *a*_1_ = −30 provides strong delayed inhibition via the direct pathway, while *a*_2_ = +15 introduces moderate excitation via the indirect pathway. This setup shows a natural balance when at rest, where inhibition stops excessive activity, but some indirect support keeps a steady rhythm. *b* _1_ and *b* _2_ values are gradually increased from zero to higher positive values to introduce varying degrees of negative saturation for the direct pathway and positive saturation for the indirect pathway. This model simulates biological damping, wherein heightened activity encounters augmented resistance.

In the hyperactive state, both routes exhibit excitatory characteristics with *a*_1_ = + 30 and *a* _2_ assigned values of + 5 or + 15, contingent upon the intensity of indirect facilitation. The nonlinear coefficients have both small positive and negative values of *b*_1_ and *b*_2_. The inhibitory state from the indirect pathway (*a*2= − 5). This imbalance dampens motor output. The nonlinear dynamics are controlled by assigning positive and negative values to both *b*_1_ and *b*_2_. Negative values of *b*_*1*_ enhance self-facilitation in the direct pathway but are counteracted in the indirect pathway, where negative *b*_2_ values result in strong suppression.

By changing the signs and sizes of these parameters in a careful way, the model looks at how neural feedback systems move between normal rhythmic patterns and unhealthy disruptions. The combination of simple feedback, complex adjustments, and timing-based design offers a flexible way to understand how the motor system works in both healthy and unhealthy situations.

This study fills a gap in research by creating a mathematical model based on VdP to explore how two types of feedback delays affect the internal resonance and oscillation patterns in complex neural systems. It investigates how delay-driven feedback and signal interactions influence system stability, enabling either oscillation persistence or disruption. These dynamics are particularly relevant to abnormal motor patterns observed in Parkinson’s disease, as discussed in previous foundational works [73–76].

A main aim is to explore the relationship between internal resonance and movement dysfunctions. We hypothesize that disrupted feedback timing exacerbates pathological rhythms and motor symptoms. The model helps examine how dual delays and nonlinear feedback induce and sustain abnormal activity. One major application is improving DBS techniques. Understanding how time delays modulate the stability of basal ganglia circuits allows better tuning of stimulation protocols to suppress pathological activity and restore healthy motor behavior [77–79].

Beyond neurology, this framework supports research into delay-governed nonlinear systems across disciplines such as physics, control engineering, and biological modeling [80–82]. It provides analytical tools to probe feedback instability and dynamic transitions in delayed systems [83–86].

Our method computes eigenvalues as a function of delay to locate Hopf bifurcation points and study transition behavior. Bifurcation and Lyapunov analyses help find changes between stable, almost repeating, and chaotic states, and they check how sensitive these states are to starting conditions.

The model also shows that when *τ*_2_ ≈ *τ*_1_, unusual beta-band oscillations (13-30 *Hz*;) can occur, which appear as resting tremor. Dysregulated timing of excitation-inhibition loops may also explain bradykinesia or dyskinesia. While the model does not fit patient-specific data, its parameters reflect physiologically plausible values, aiming to reproduce Parkinsonian oscillatory behavior qualitatively.

Additionally, our results support the hypothesis that clinical symptoms originate from disrupted signal timing in basal ganglia circuits. Delays in the indirect pathway keep the motor cortex excessively primed, causing resting tremor. Conversely, delayed initiation signals hinder motor activation, producing bradykinesia followed by rebound hyperactivity. As these delays lengthen with disease progression, symptoms intensify. In extreme cases, long delays in the direct pathway may induce chaos, offering an explanation for the disordered movements in Huntington’s disease. This paper presents a unified framework for delay-driven movement pathologies.

Building on these insights, our focus centers on two key directions:

- Studying symptom progression with increasing delay: We assess how escalating delays in motor circuits influence stability and movement.
- Characterizing resting-state resonance: We investigate how time-delay interactions generate beta oscillations linked to tremor.

Through this targeted exploration, we aim to deepen understanding of delay-driven motor dysfunctions and inform improved therapeutic interventions.

## 2. Analyzing Resonance Dynamics

This study looks at how longer delays in motor feedback loops affect movement by examining the delayed VdP oscillator in two stable conditions: one where there is no movement (*x*^*^ = 0) and one where movement is happening 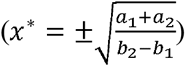. This provides a clear way to see how differences in time delays can lead to harmful changes in oscillations. Hopf bifurcation analysis is used to identify when self-sustained oscillations begin due to these delayed interactions. These represent the inactive resting state (x^*^ = 0) and the active movement-generating state 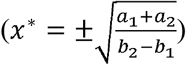, offering a simplified yet powerful framework to track how time delay differences influence pathological oscillatory transitions [83].

Hopf bifurcation analysis is used to detect the onset of self-sustained oscillations as a function of delayed interactions. The zero-equilibrium case shows how frequency changes when the system is at rest, while the non-zero equilibrium explains how movement states become unstable when the timing of excitation and inhibition is changed. The key of relationships is captured by the equations:

### For the resting state, oscillations emerge when

For the zero-equilibrium point:

Real part:

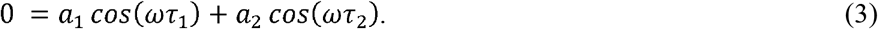

Imaginary part:

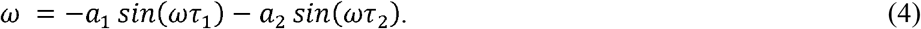

Combined equation (squaring and adding):

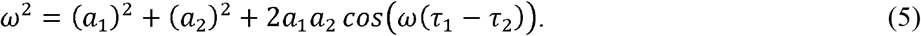

### For the movement-active state

Real part:

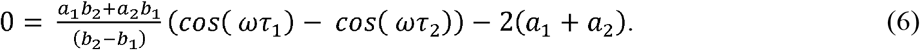

Imaginary part:

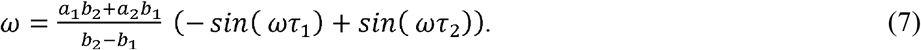

Combined equation (squaring and adding):

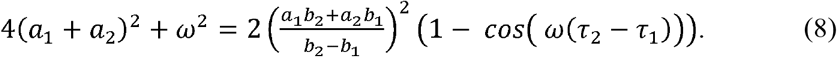

These expressions demonstrate how increasing delay differences—especially in the indirect pathway—reshape the natural frequencies of the system. The shift to unhealthy rhythms relies not just on how much delay there is but also on how excitation (*a*_2_) and inhibition (*a* _1_) work together, which is shown by their effects on the rhythm’s behavior.

#### 2.1 Internal Resonance Conditions

Internal resonance happens when the natural frequencies of a system are related by a simple number ratio, which lets energy move between pathways the system can oscillate. In biological systems such as neural networks that rely on delayed feedback, understanding these conditions is essential for predicting and controlling oscillatory disturbances.

The governing equation for the first pathway with time delay is expressed as:

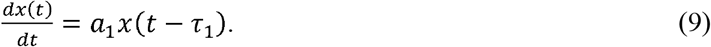

The characteristic equation obtained from the linear stability analysis leads to the following conditions:

Real Part:

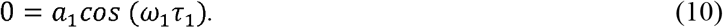

Imaginary Part:

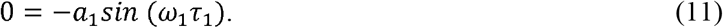

Therefore, the natural frequency can be determined as:

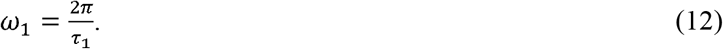

Equating the magnitude of the oscillatory terms, to obtain

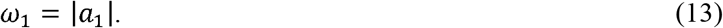

Hence, the constraint on *a*_1_ has the form

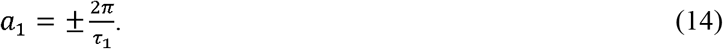

Similarly, for the second pathway, one obtains

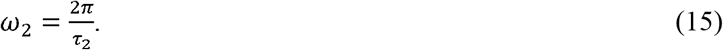

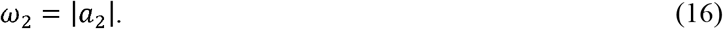

Thus:

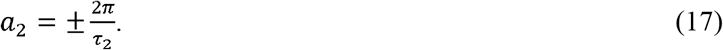

It is significant to mention that the internal resonance occurs when the ratio of the natural frequencies is a rational number:

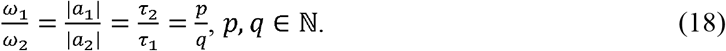

The superharmonic resonance condition arises when one frequency is a multiple of another:

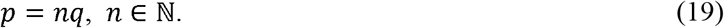

The subharmonic resonance occurs when one frequency is a submultiple of another:

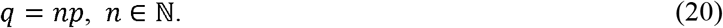

To accurately calculate the final resonant frequency, the resonance conditions are replaced in oscillation equations (5) and (8) as follows:

For the zero equilibrium point:

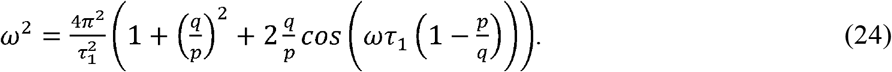

For the non-zero equilibrium point:

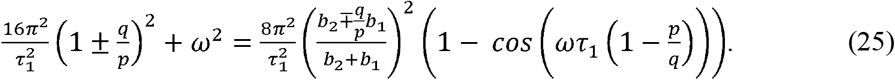

Examining the impact of the internal delay structure on the angular frequency response reveals that resonance behavior in delayed feedback systems is highly sensitive to both the delay ratio and the configuration of linear and nonlinear feedback. The system’s ability to keep, boost, or reduce oscillations depends on a complicated relationship between the direct and indirect pathways, where their delays and gain settings affect the resonance ratio 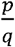. The following analysis evaluates this relationship across three systematically varied regimes.

Figure 1 shows how the angular frequency *ω* changes based on the resonance ratio 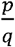, calculated when the system is at rest and the time delay in the direct pathway is set to 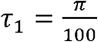. The resulting frequency profile exhibits dense, irregular fluctuations in *ω*, particularly in the low-to-intermediate range of 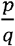. This strong nonlinear behavior indicates that even tiny differences in how long the pathways take can cause the system to break into different patterns and become unstable. The system does not settle into a frequency-locked regime, indicating a high sensitivity to delay ratios even under resting conditions. Even without nonlinear changes, the natural differences in delay are enough to cause unpredictable shifts between different oscillation patterns (Fig. 1).

**Fig. 1:**
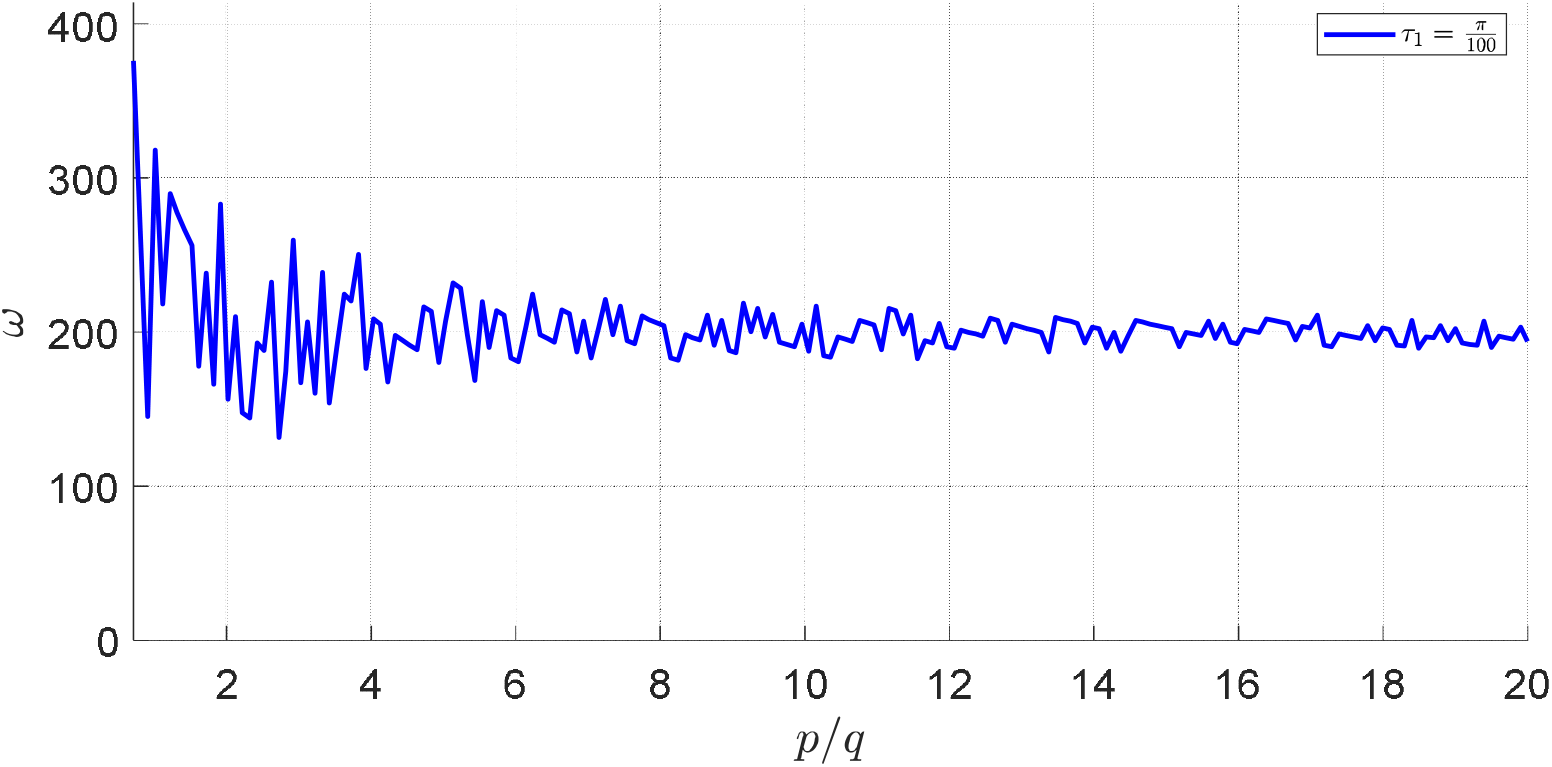
Variation of the modified frequency *ω* vs. resonance ratio 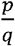 in (24), with 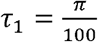.

Figure 2 demonstrates how varying degrees of nonlinear saturation affect oscillatory stability. The nonlinear feedback terms vary across three configurations: one with negative saturation (*b*_1_ = −, *b*_2_ = +0.10) and two with mixed saturation (*b*_1_ = −, *b*_2_ = +0.10) and *b*_1_ = +0.05, *b*_2_= +0.30). In all cases, the time delay is held at 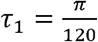. The graphs show a clear change from unstable resonance at low 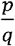 to regular frequency steps as 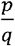 increases. This staircase pattern reflects a classical signature of frequency locking, where the system entrains to specific delay ratios. Importantly, systems with stronger nonlinear facilitation start to show oscillations later and have wider areas of suppression around 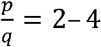, followed by a quicker return of resonance. These behaviors are similar to conditions like dyskinesia, where there are sudden bursts of activity followed by quiet periods because of unstable internal feedback (Fig. 2).

**Fig. 2:**
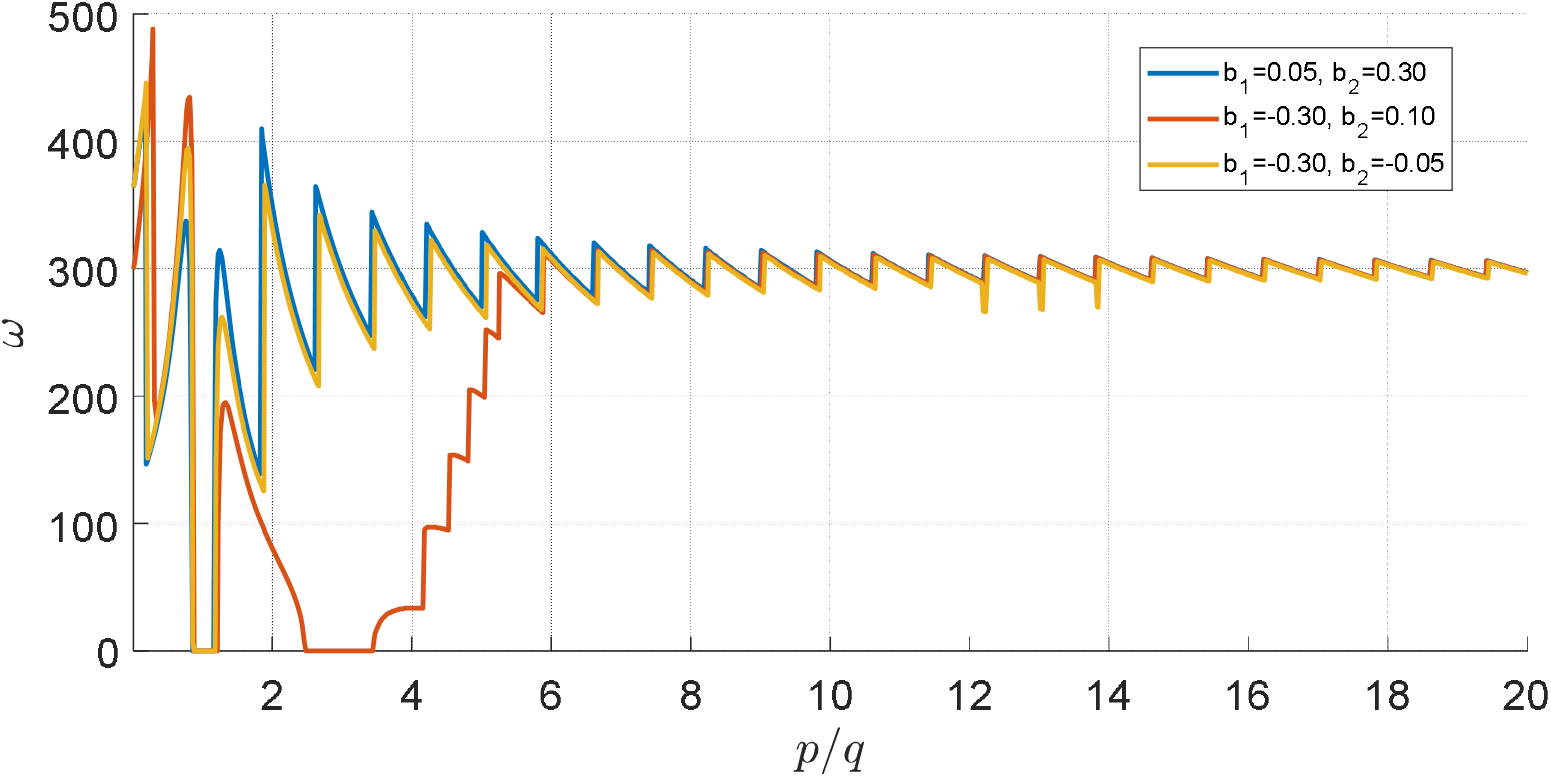
Variation of the modified frequency *ω* vs. positive resonance ratio 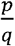 in (25), with 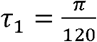.

Figure 3 looks at how resonance changes under three different sets of parameters, each showing various levels of nonlinear saturation. When *a*_1_= +30 *a*_2_ = +5, *b*_1_ = +0.05 and *b*_2_ = +0.30, there are clear resonance peaks at low *p*/*q* ratios and a slow decline, showing a strong excitatory drive that is countered by indirect suppression. When *b*_1_ = −0.30, and *b*_2_ = +30, it generates sharp and repeated resonance spikes, suggesting persistent excitation and b_2_ = −0.05. The result is a suppression of early resonance, followed by a and enhanced sensitivity to timing mismatches. In the third configuration, when *a*_2_ = +15, b_1_ = −0.30 and b_1_ = −0.05. progressive build-up at higher delay ratios. This behavior shows a way to keep things stable at short delays, but it becomes unstable at longer feedback loops, which might explain the change from stable to problematic resonances seen in motor disorders like Parkinson’s disease (Fig. 3).

**Fig. 3:**
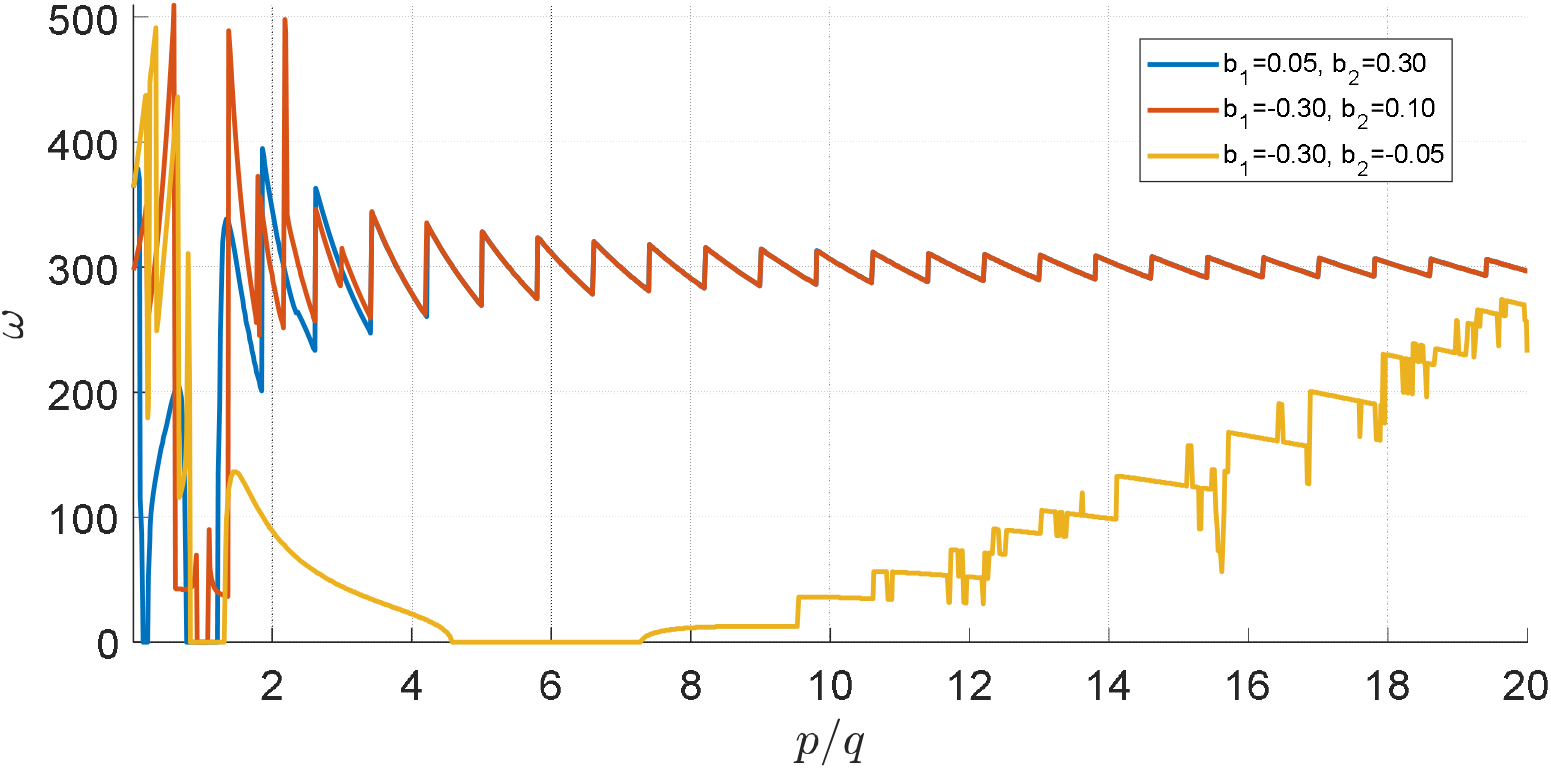
Variation of the modified frequency *ω* vs. negative resonance ratio 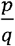 in (25), with 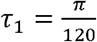.

Together, the aforementioned figures show that delay-dependent internal resonance is not a constant feature of the system but a flexible process influenced by how excitation, inhibition, and nonlinear saturation are arranged. In particular, the ratio 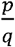 acts as a control parameter that determines whether oscillations are enhanced, stabilized, or fully suppressed. Moreover, the nonlinear gains *b*_1_ and *b*_2_ introduce asymmetry between pathways, allowing fine-tuned modulation of amplitude and frequency. This feature has strong implications for motor control and for the design of adaptive neuromodulation systems, where real-time feedback could be used to identify and disrupt harmful resonance patterns by manipulating delay structure or saturation parameters.

## 3. Numerical Simulations and Bifurcation Analysis

To systematically examine how variations in feedback delays influence the qualitative behavior of the system, this section presents a set of numerical simulations and bifurcation analyses. By integrating delayed nonlinear dynamics with diagnostic tools such as bifurcation diagrams and Lyapunov exponent calculations, we aim to reveal the rich landscape of possible dynamical regimes—including steady states, oscillations, and chaos—that arise under biologically plausible conditions. Although equations (24) and (25) demonstrate that feedback delays can give rise to oscillatory behavior, they are often based on simplified assumptions such as perfect resonance or symmetry. In contrast, noise, variability, and adaptive constraints influence the near-resonant regimes in which biological systems operate. To bridge this gap between theory and biology, we conducted numerical simulations to explore the system’s bifurcation landscape as a function of the secondary delay *τ*_2_, while keeping the primary delay *τ*_1_fixed. These simulations used realistic values that show how excitatory and inhibitory loops interact in cortical-basal ganglia circuits.

The core of the analysis combines two complementary diagnostic tools:

- Bifurcation diagrams generated from simulations of a delayed VdP-like oscillator.
- Estimation of the largest Lyapunov exponent (LLE) to quantify dynamic stability.

Bifurcation diagrams illustrate the asymptotic values of the system state *x*(*t*) across varying *τ*_2_. These figures enable visual identification of transitions between steady-state, periodic, multi-periodic, and chaotic dynamics.

To better understand these transitions, we calculated LLE, which shows how nearby paths in phase space either move apart or come together on average:

- *λ*< 0: stable periodic or fixed-point attractors.
- *λ* = 0: quasi-periodic behavior or critical bifurcation threshold.
- *λ* > 0: chaotic and unstable dynamics.

This measure provides a quantitative framework for identifying changes in system behavior under varying delay conditions.

### 3.1 Finite-Difference Estimation

In the preliminary analysis (Figs. 4–6), a rapid approximation of the Lyapunov exponent was obtained by computing the logarithmic derivative of the solution trajectory over the final portion of each simulation. For each value of *τ*_2_, the slope was estimated using the last 5% of the time series *x*(*t*) that was extracted after numerically solving the DDE using MATLAB’s dde23. The approximate derivative 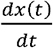 was estimated using finite differences:

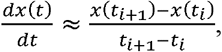

subsequently, Lyapunov exponent was approximated as:

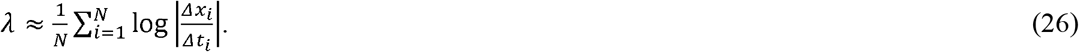

**Fig. 4:**
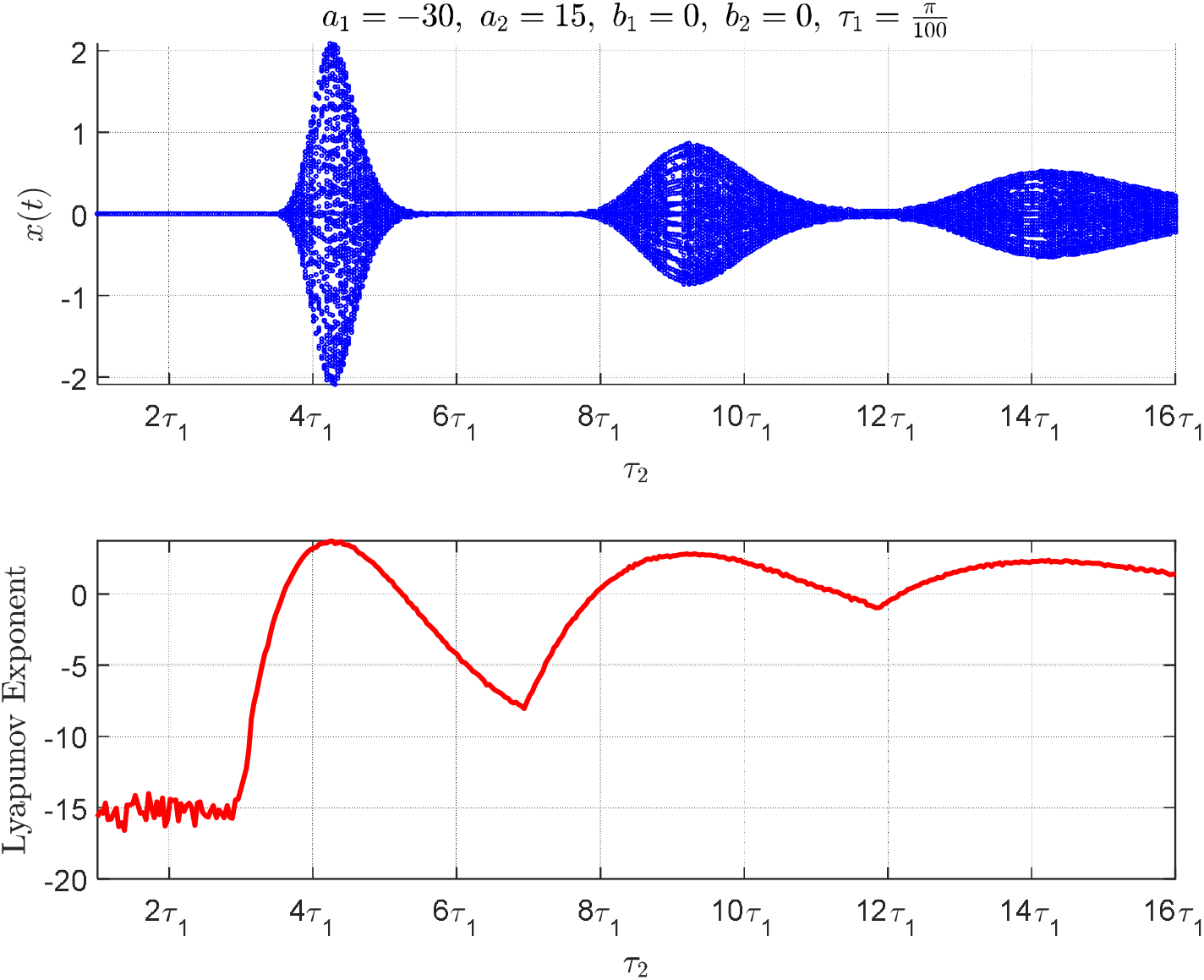
Bifurcation diagram and Lyapunov exponents from (2) without nonlinear interaction in the resting state, using the final 5% of a 10 − *second* simulation.

This derivative-based logarithmic mean offers a rapid overview of the system’s divergence behavior.

#### Advantages

- Computationally efficient and easy to implement.
- Requires only the original system—no augmentation needed.

#### Limitations

- Highly sensitive to noise and discretization errors in estimating derivatives.
- Fails to robustly differentiate between weak chaos and quasi-periodic dynamics.
- Underperforms in systems with strong nonlinear damping, where small amplitude fluctuations mask underlying instability.

As revealed in our simulations, this method produced ambiguous or misleading estimates in regimes with nonlinear saturation (e.g., positive/negative values of *b*_1_, *b*_2_), especially when oscillations remained bounded but complex.

This method serves as a qualitative screening tool for detecting bifurcation points. However, it lacks sensitivity in identifying weak chaos or high-frequency modulations typical of neurophysiological rhythms.

### 3.2 Benettin’s Variational Algorithm

To make it more accurate—especially for spotting complicated changes like resonance breakdown or deterministic chaos—a changed version of the Benettin algorithm was used. This approach involves solving an augmented system of delay differential equations that includes both the nominal trajectory *x*(*t*) and a small perturbation *δ x*(*t*). The system governs the evolution of perturbation.

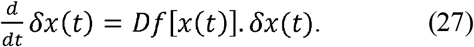

Here, *Df* denotes the Jacobian of the nonlinear DDE system, manually computed for the nonlinear saturation terms. The augmented system in the simulation integrates both the base signal 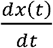 and the perturbed signal 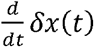.

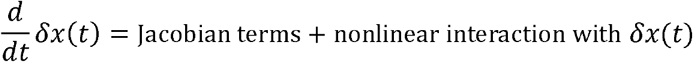

Perturbations are re-normalized periodically to avoid numerical blow-up. The final LLE is calculated by averaging the logarithmic divergence of the perturbation over time:

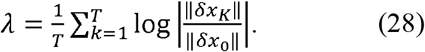

#### Advantages

- Accurately detects deterministic chaos, high-order bifurcations, and resonance collapse.
- Captures effects of nonlinear damping and saturation that the finite-difference method misses.
- Maintains numerical stability via explicit re-normalization.

#### Limitations

- Computationally more expensive.
- Requires symbolic or numerical derivation of Jacobian terms.
- Sensitive to the correct initialization of perturbation magnitude *δx*_0_

While the finite-difference approach was suitable as a preliminary bifurcation screening tool, it failed to resolve fine-grained instabilities induced by the system’s nonlinear structure. Particularly, it could not distinguish mild instability from quasi-periodicity, nor could it track the sensitivity arising from negative nonlinear damping terms that are biologically plausible in motor control dysfunctions.

In contrast, the Benettin variational method was found indispensable for analyzing systems with strong or sign-changing saturation terms, where the response surface becomes highly sensitive to initial perturbations. This sensitivity is a hallmark of systems approaching chaotic attractors or exhibiting transient resonance.

Figure 4 shows absence of nonlinear saturation (*b*_1_ = *b*_2_ = 0), the bifurcation diagram reveals a gradual transition from periodic to multi-periodic oscillations as *τ*_2_ increases. The Lyapunov exponent remains negative until *τ*_*2*_*≈* 4.1*τ*_1_, indicating stable behavior. A bifurcation occurs at *τ*_*2*_*≈* 4.2*τ*;_1_, initiating a cascade of increasingly complex dynamics. This setup is like the early signs of Parkinson’s tremor, where little resistance and strong positive feedback cause uneven shaking (Fig. 4).

Figure 5 shows moderate nonlinear feedback (*b*_1_ = + 0.30 *b*_2_ = + 0.15), with mild nonlinear damping, the system exhibits more distinct bifurcation branches. Oscillations remain suppressed until *τ*_*2*_*≈* 4.2*τ*_*1*_, after which instability begins around *τ*_*2*_*≈* 4.2*τ*_*1*_, as the LLE approaches zero. This behavior is like how inhibitory feedback systems, like GABAergic transmission, work by postponing harmful oscillations while still permitting brief periods of resonance. Such dynamics may represent transitional hyperkinetic motor states where short-lived instabilities cause intermittent uncontrolled movements (Fig. 5).

**Fig. 5:**
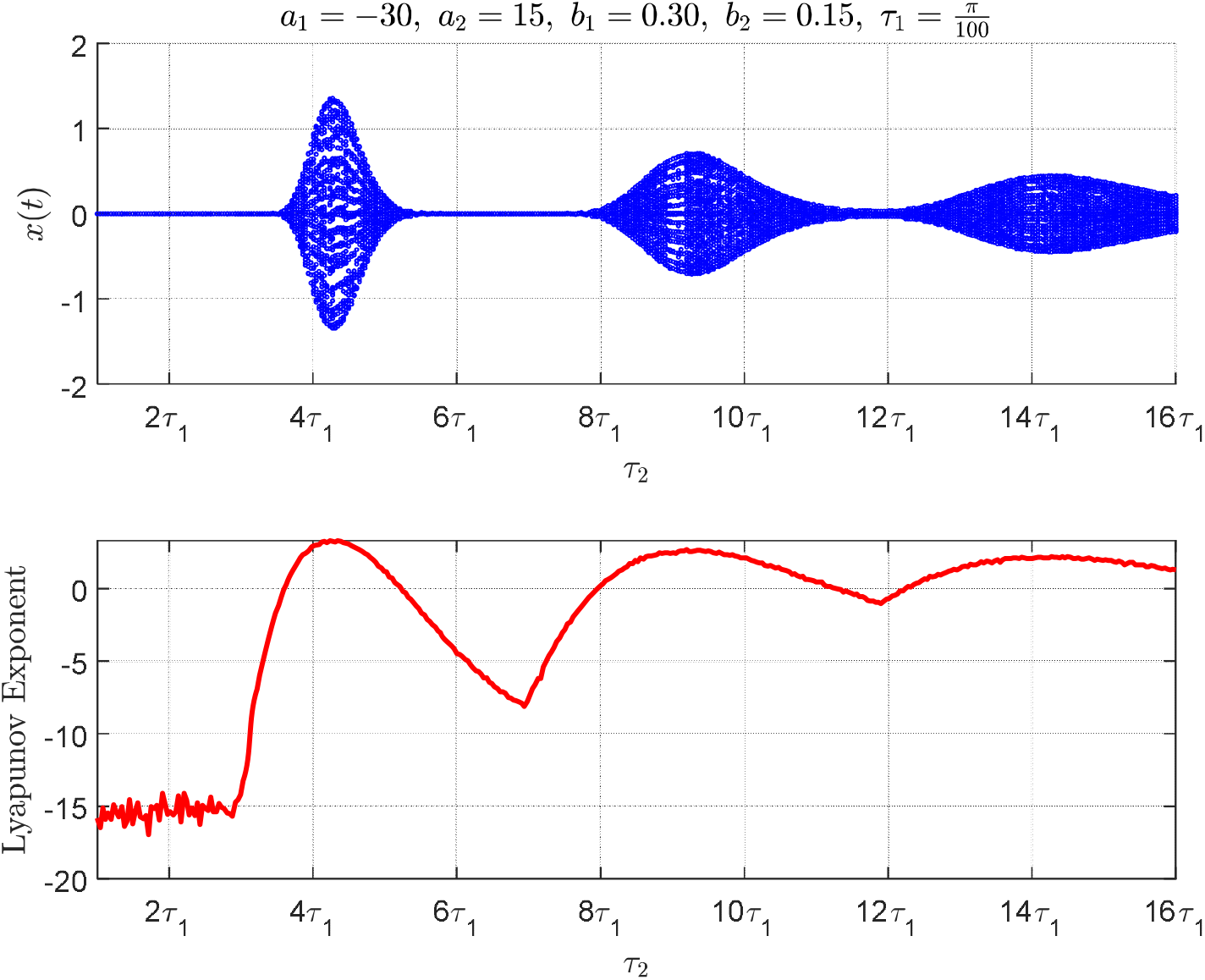
Bifurcation diagram and Lyapunov exponents from (2) with moderate nonlinear interaction in the resting state, using the final 5% of a 10 − *second* simulation.

Figure 6 shows strong nonlinear saturation (*b*_1_ = + 1.80, *b*_2_ *b*_2_ = + 0.90) under strong nonlinear saturation, the system becomes sensitive to mismatched delays. The first bifurcation appears early, near *τ*_2_ *≈* 3.2 *τ*_1_, and the LLE rises above zero, signaling the onset of deterministic chaos (Fig. 6). These behaviors are like what happens in conditions like deep cortical dysrhythmia or tremors in Parkinson’s disease, where timing errors in feedback systems lead to chaotic movement signals.

**Fig. 6:**
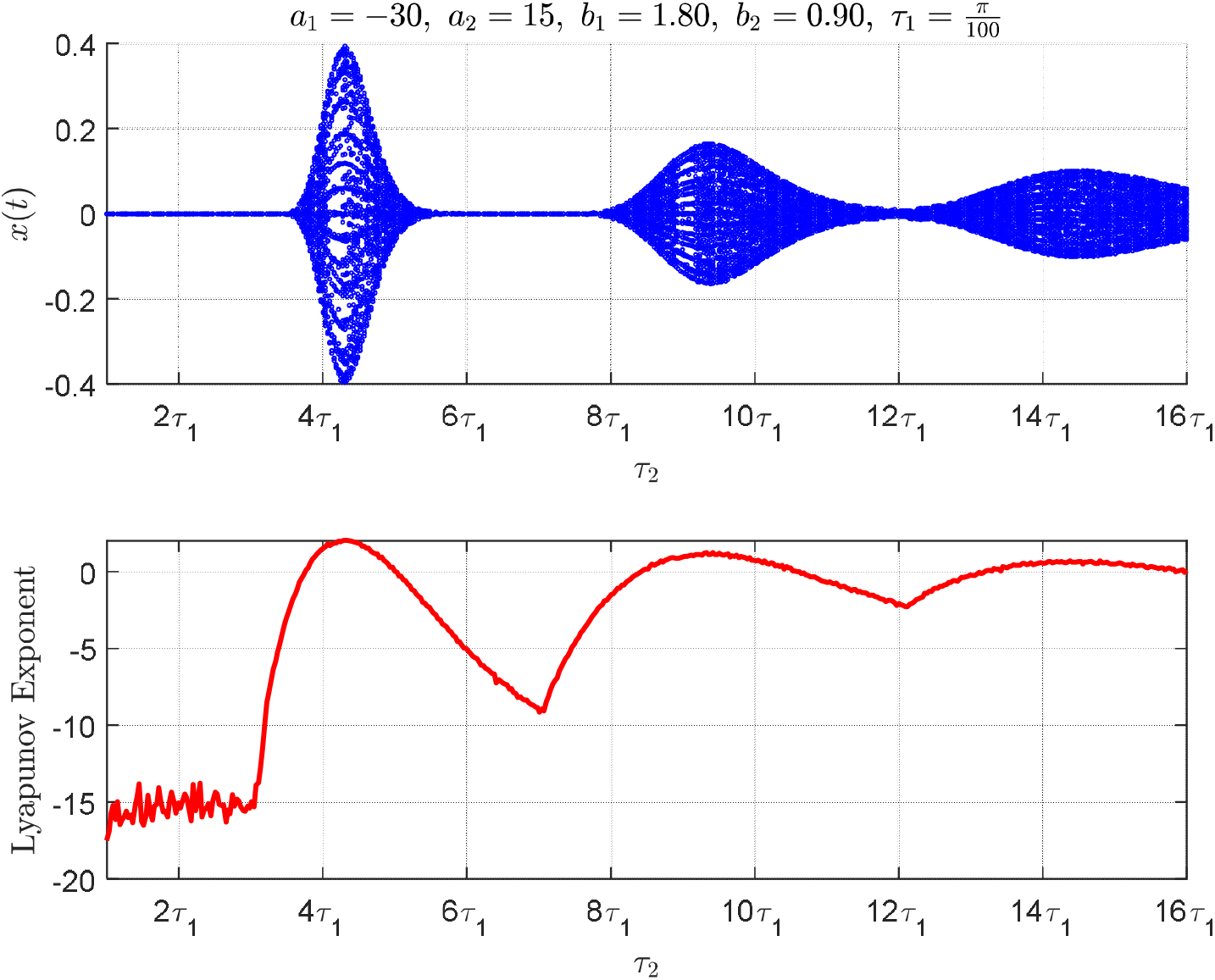
Bifurcation diagram and Lyapunov exponents from (2) with strong nonlinear interaction in the resting state, using the final 5% of a 10 − *second* simulation.

Without saturation, the system doesn’t have a built-in way to control too much excitement, which causes ongoing oscillations—like the tremors seen in Parkinson’s disease, where the ability to inhibit is weakened and the excitement of neural circuits goes up (Fig. 4). With moderate saturation, the system exhibits a greater tolerance to increased delay and maintains stability for a longer period. This behavior resembles the natural regulation of movement via mechanisms such as GABAergic neurotransmission and intact dopaminergic modulation (Fig. 5). The eventual instability may reflect transitional states such as treatment-induced involuntary movements. With strong saturation, the system becomes hypersensitive to small temporal changes and rapidly degenerates into a chaotic state. This stage shows the strong connection between brain waves in the beta band seen in patients with advanced Parkinson’s disease, where too much coordination between brain circuits causes a loss of movement control and symptoms like being unable to move (Fig. 6). Nonlinear saturation is a crucial component of the regulation of neuromotor dynamics. If there is not enough nonlinear saturation or none at all, the stability range becomes limited, but too much saturation makes the network less flexible and can cause it to fail.

Figure 7 shows a setup with positive nonlinearity and increased excitability, using the values *a*_1_ = + 30, *a*_2_ = + 5, *b*_1_ = 0.05, *b*_2_ = + 0.30 and 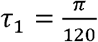. The time series of *x*(*t*) reveals an abrupt bifurcation around *τ*_2_ ≈ 6*τ*_1_, followed by large-amplitude oscillations that stabilize into a periodic plateau (Fig. 7). This behavior shows how the hyperdirect pathway helps reduce too much excitement caused by nonlinear feedback interactions. In Figure 8, the negative value of *b*_1_ introduces negative saturation within the direct pathway, effectively enhancing excitatory gain. However, this configuration results in mild suppression of activity, leading to dampened oscillatory behavior and simple stability (Fig. 8). Figure 9 presents a balanced configuration of negative nonlinearities with *a*_1_ = + 30, *a*_1_ = + 15, *a*_1_ = − 30 and *a*_1_ = − 0.05. Despite the lack of traditional damping, the oscillation amplitude remains confined within a narrow range, and LLE remains negative and stable, possibly indicative of bounded low-dimensional chaos (Fig. 9). In Figure 10, further amplification of positive saturation from the indirect pathway (*b*_2_ = −0.15) destabilizes the system over time. This results in growing oscillatory amplitudes and frequency jittering. The corresponding LLE starts from a deeply negative value and gradually approaches to zero, signifying a transition toward instability or multi-stability, possibly approaching a chaotic attractor landscape (Fig. 10). Figure 11 examines the influence of the simulation time window by comparing outputs over *t*_*final*_ and *t*_*final*_ = 1 s using parameters *a*_1_ = + 30, *a*_1_ = + 5, *a*_1_ = − 30 and *a*_1_ = +0.10. The extended simulation clarifies how prolonged response delays in the indirect pathway can lead to system dynamics that either undershoot or overshoot the equilibrium threshold. This behavior maps to clinical phenomena such as bradykinesia or motor hyperactivity in Parkinsonian states, reflecting the dual roles of inhibition and disinhibition in motor control circuits (Fig. 11). Altogether, these results underscore the critical role of nonlinear saturation parameters 1 and 2 in shaping the dynamic behavior of delayed feedback neural systems. The sign and magnitude of these parameters dictate whether the system evolves toward periodic, chaotic, or stable states. Furthermore, transitions between these regimes are tightly coupled with *τ*_2_, which models long-loop cortico-basal ganglia interactions. These findings suggest that saturation-driven modulations of neural feedback can serve as measurable indicators of motor dysfunction and may offer actionable targets for therapeutic intervention via pharmacological or neuromodulatory strategies.

**Fig. 7:**
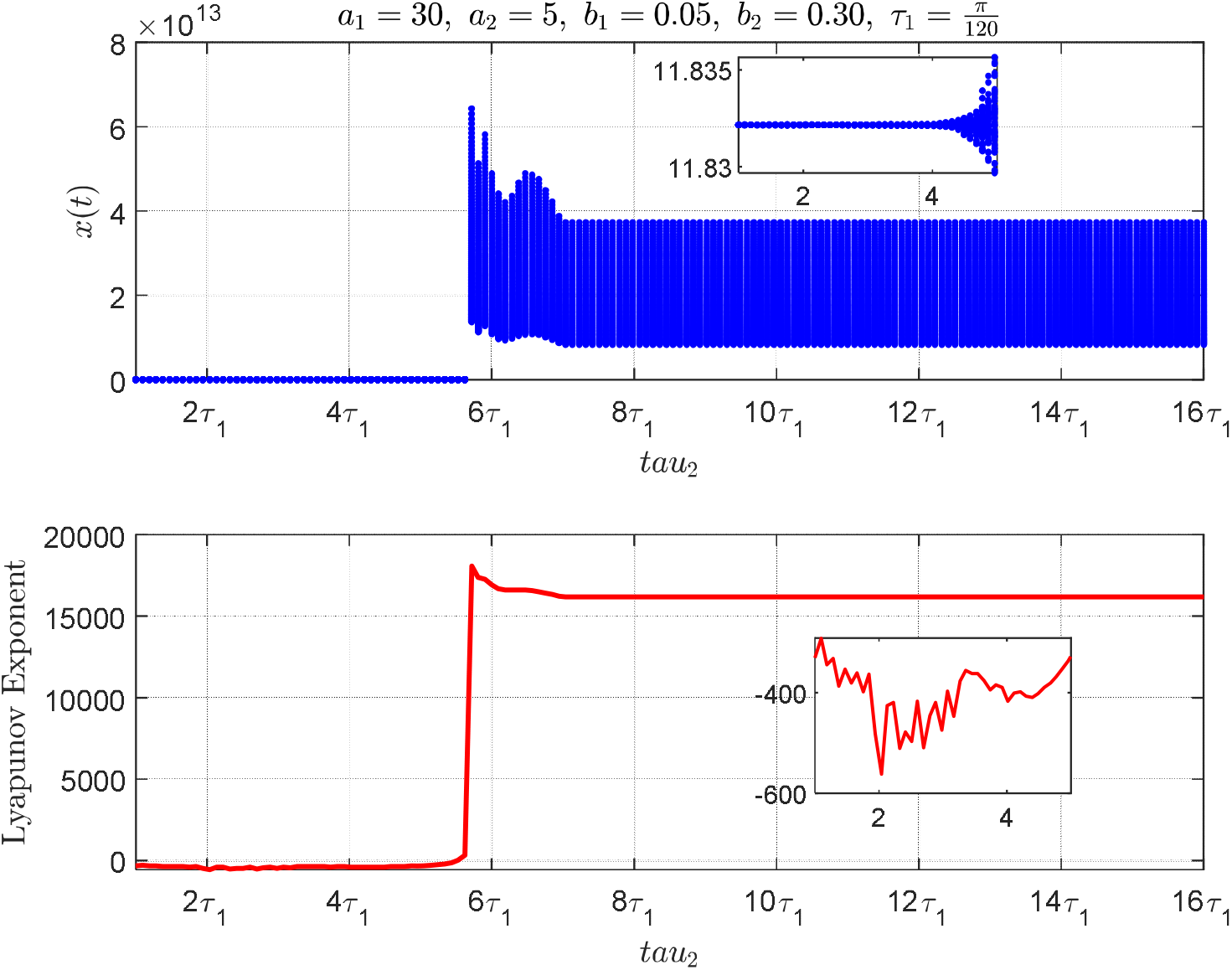
Under strong excitatory drive, a critical delay triggers large-amplitude oscillations and instability, as seen in the final 5% of a 5 − *second* simulation.

**Fig. 8:**
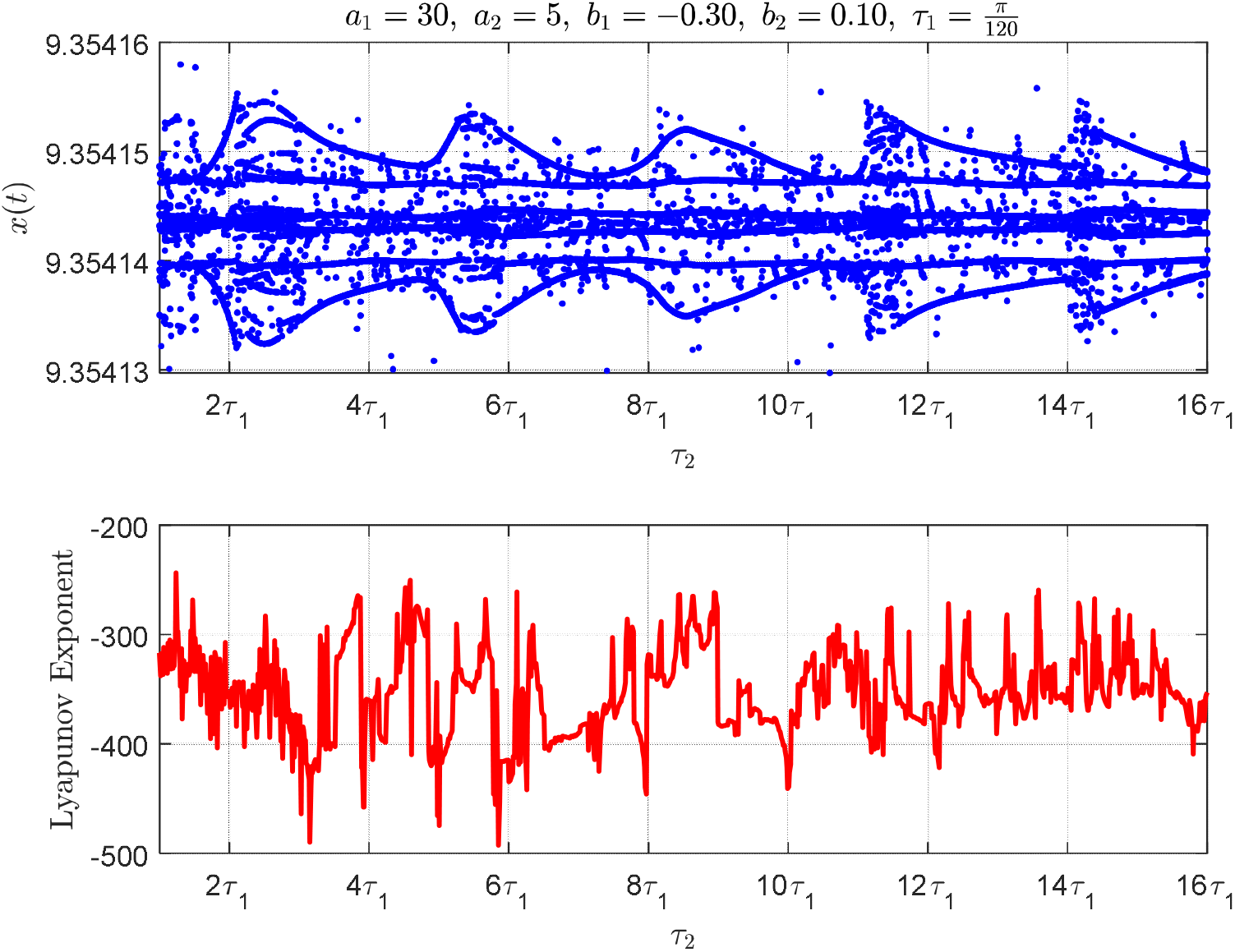
Negative saturation, the system maintains bounded oscillations and negative Lyapunov exponents across delays, confirming stability in the final 5% of a 5 − *second* simulation.

**Fig. 9:**
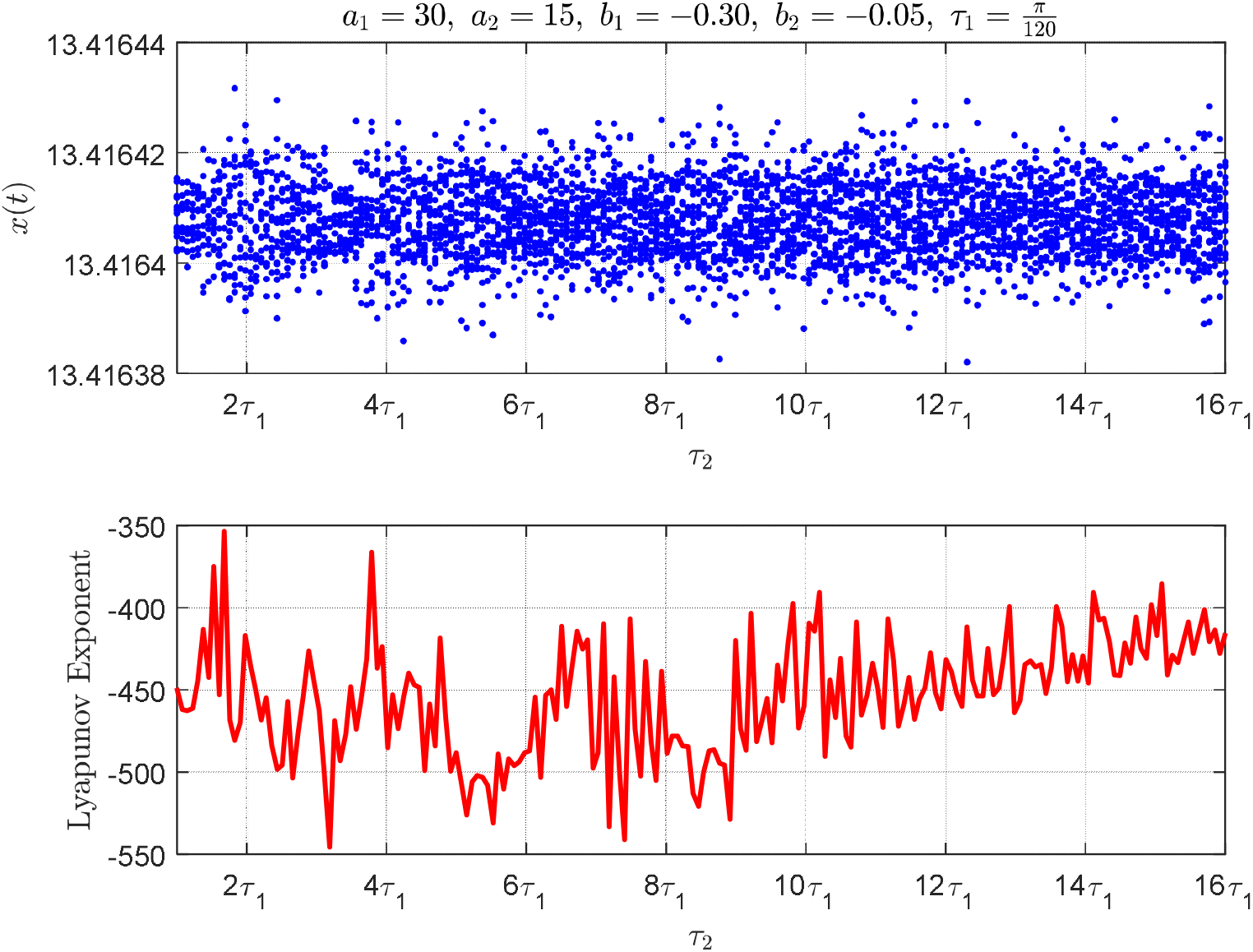
Balanced negative direct and high positive indirect pathway saturation, yields minimal variation in state trajectories, during the final 5% of a 5 − *second* simulation.

**Fig. 10:**
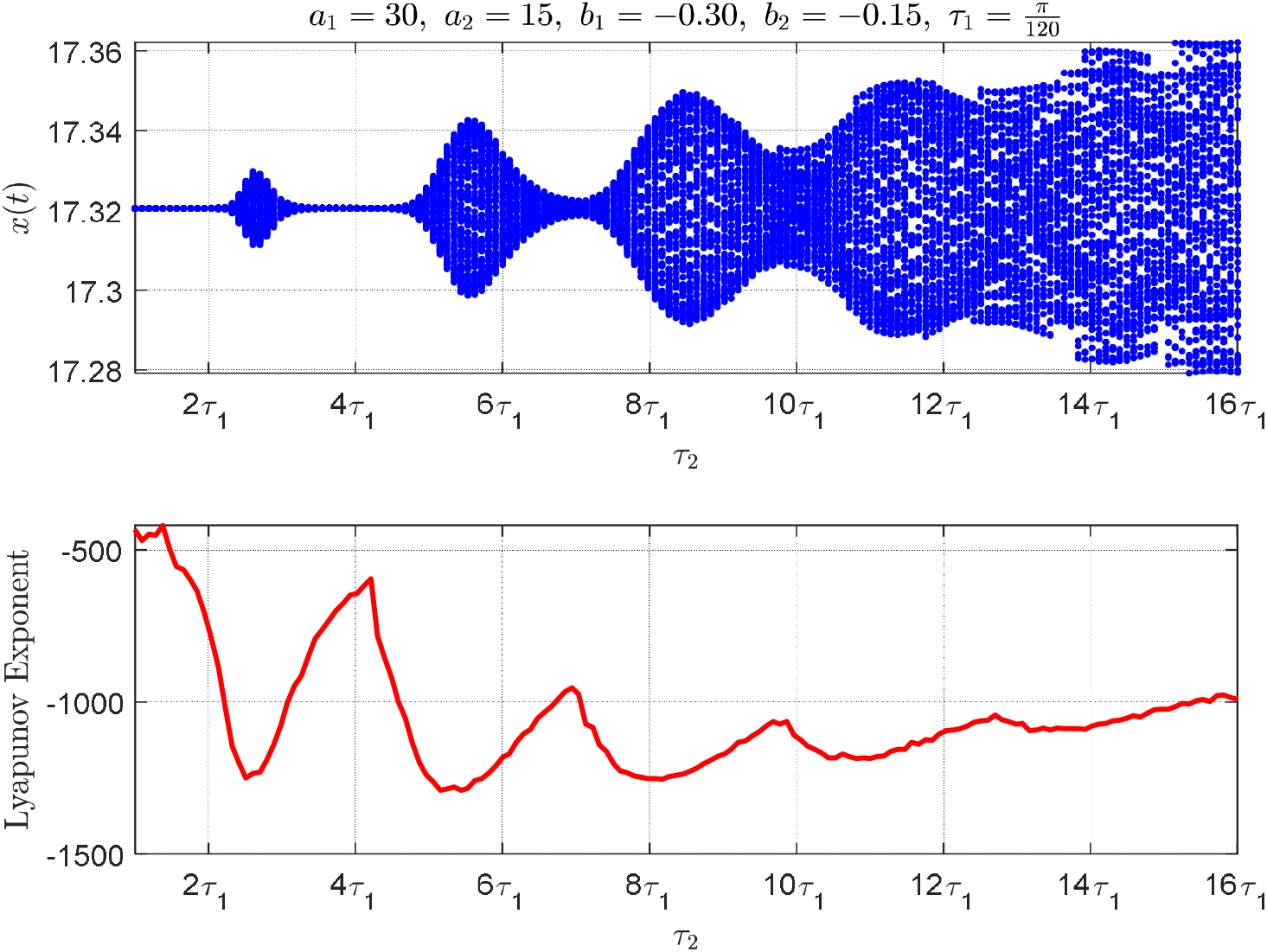
Enhanced indirect excitation leads to growing amplitude modulation and burst, destabilization in the final 5% of a 5 − *second* simulation.

**Fig. 11:**
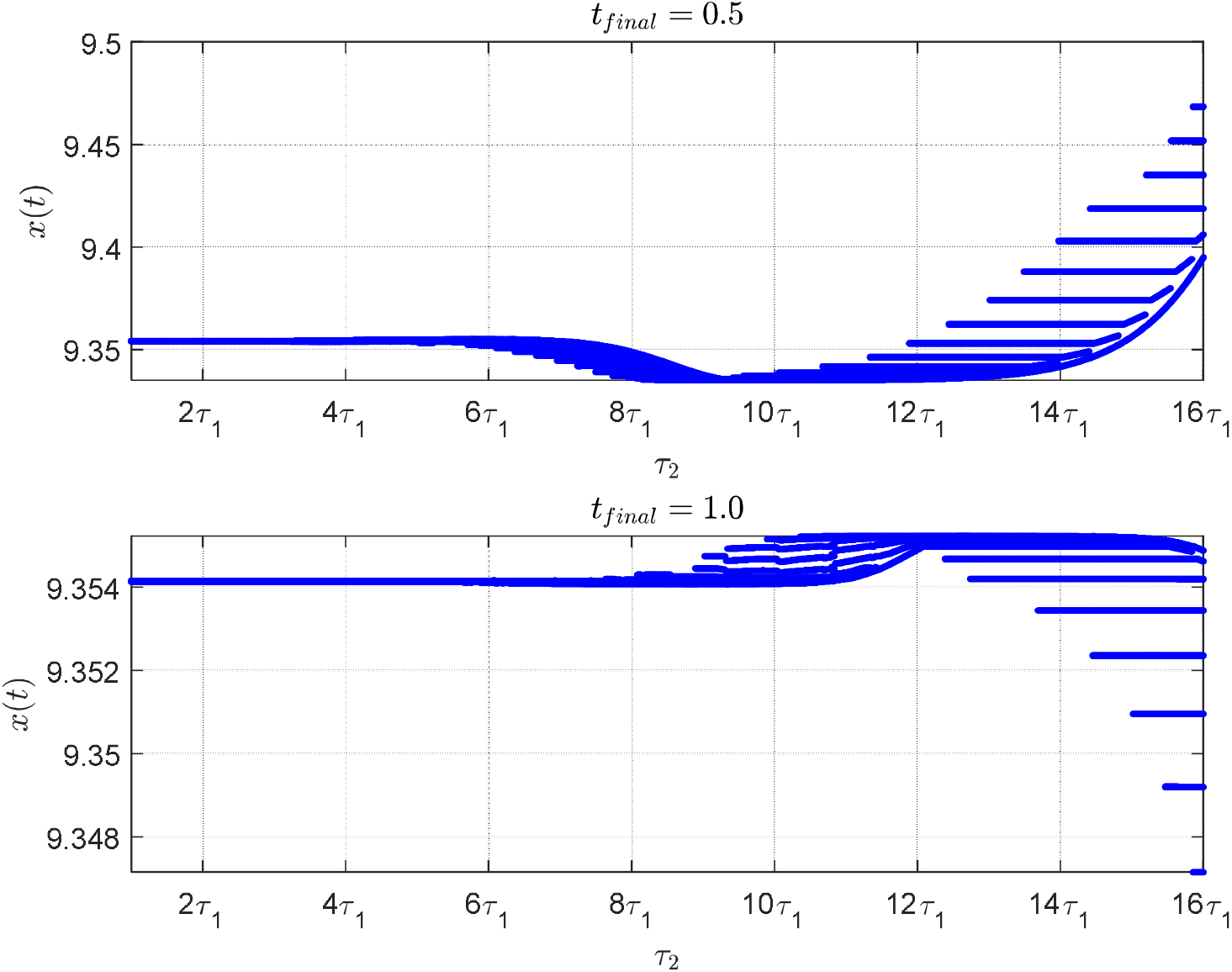
Time-dependent bifurcation patterns under moderate indirect excitation. At short duration 0.5 s and 1.0 s. Data reflects the final 5% of each simulation.

Figure 12 illustrates how a strong negative saturation in the indirect pathway, combined with a moderate increase in the direct pathway 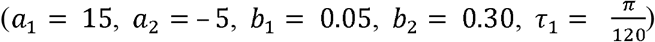, impacts the system. The system initially remains near equilibrium until a bifurcation emerges sharply at *τ*_2_ ≈ 6*τ*_1_. This bifurcation causes a big increase in size, and the oscillations change to a very high range, indicating a numerical divergence (Fig. 12).

**Fig. 12:**
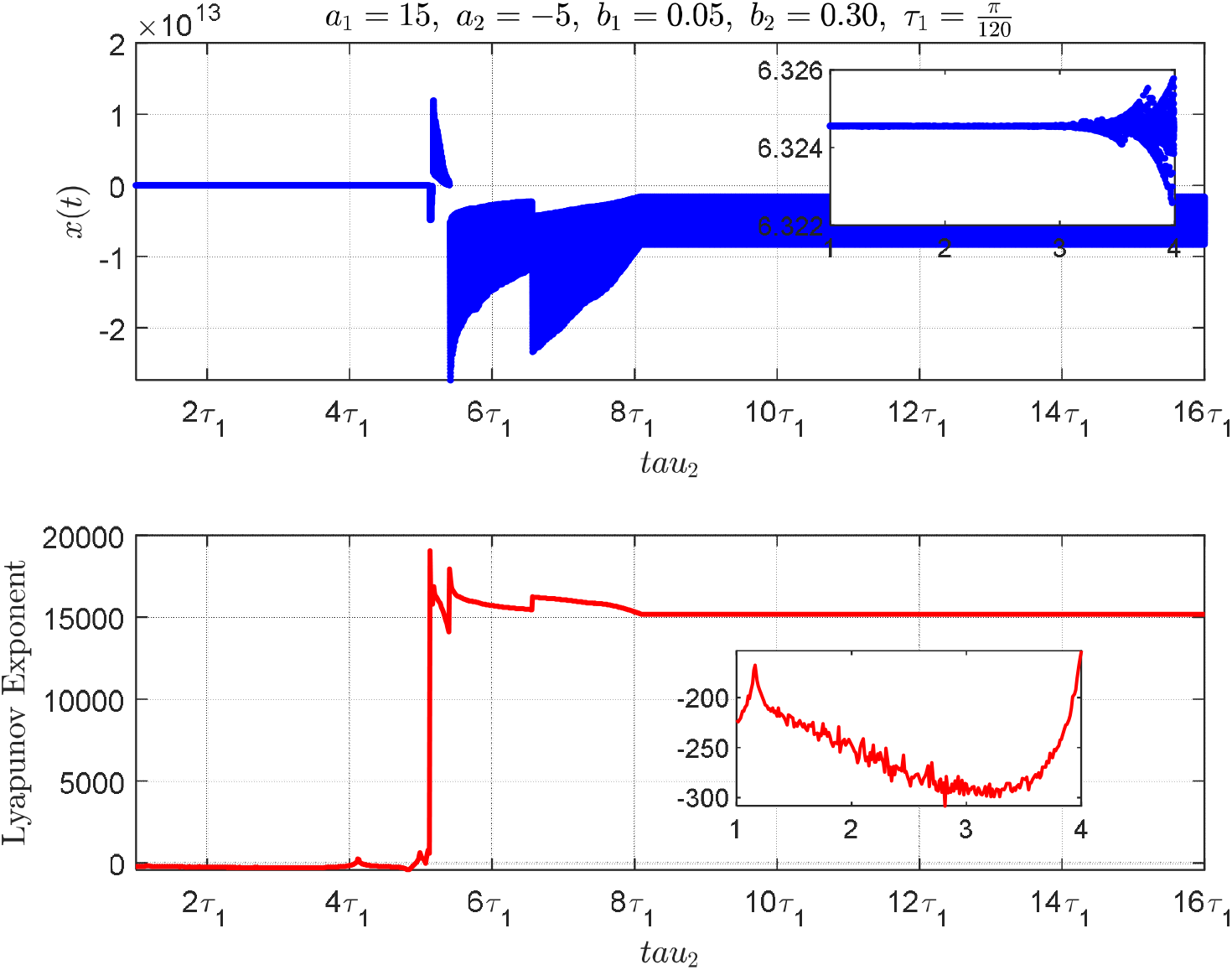
Positive saturation in the direct pathway and strong indirect feedback, the system exhibits rapid instability beyond a delay threshold. Final 5% *of a* 5 − *second* simulation.

Figure 13 shows a strong decrease in the direct pathway along with a moderate increase in the indirect pathway (*b*_1_ = − 0.30, *b*_2_ = − 0.10). Here, nonlinear facilitation is present across both feedback loops, suppressing amplitude while preserving variability. The trajectory of x(*t*) remains confined within narrow fluctuation margins, and the LLE stays negative but shows gradual ascent. This pattern indicates weakly chaotic dynamics within a stable envelope (Fig. 13).

**Fig. 13:**
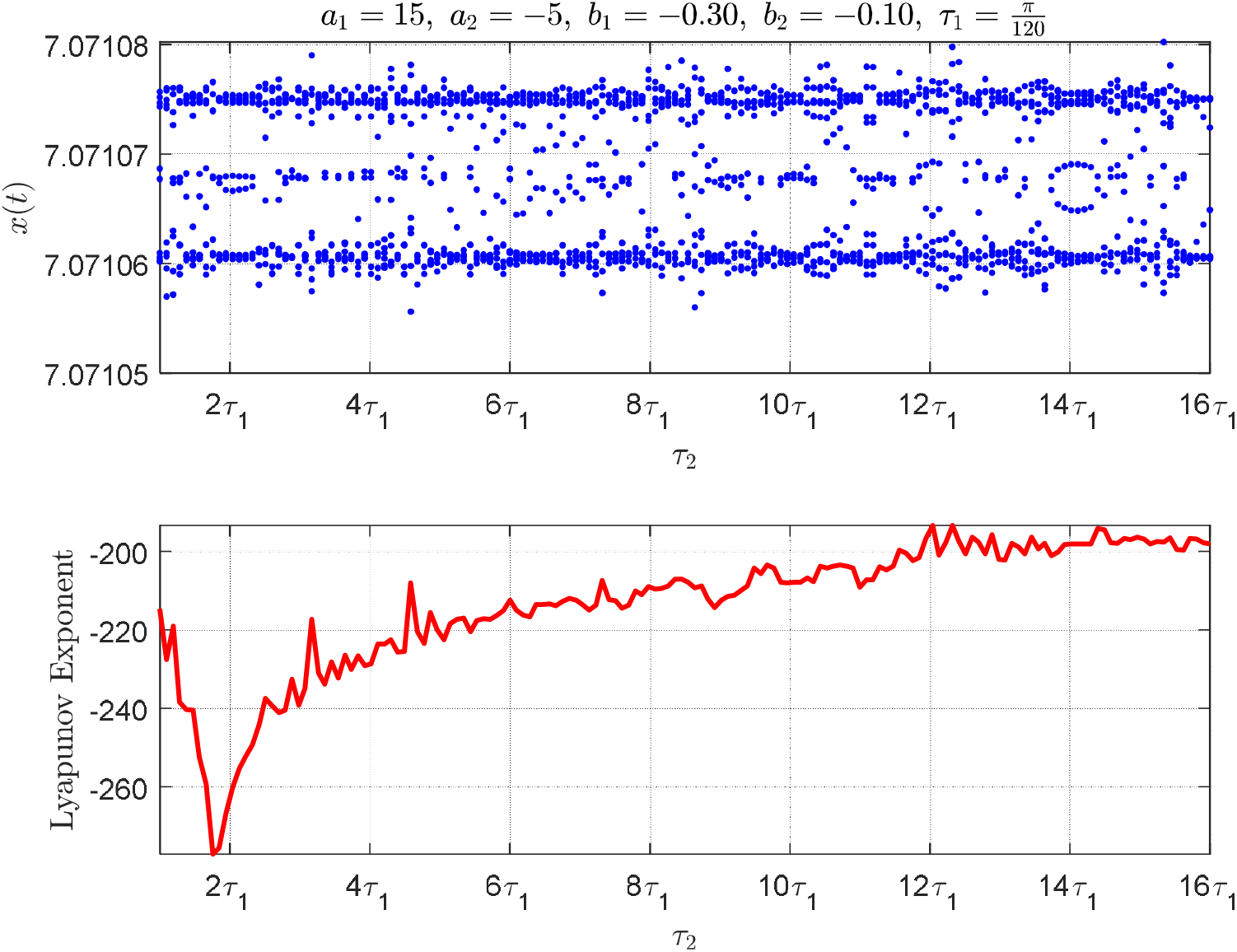
With negative saturation in the direct pathway, state trajectories remain confined and decay slowly. Final 5% *of a* 5 − *second* simulation.

In Figure 14, features moderate negative saturation in both pathways (*b*_1_ = − 0.15 b_2_ = 0.05). The dynamics show a piecewise constant trajectory, with abrupt state transitions and long persistence in discrete values. The LLE reflects these discontinuities as sharp peaks, suggesting a form of bistability or metastability (Fig. 14). The positive spikes in the Lyapunov curve support this claim. This behavior could represent moments when a person suddenly stops moving or has trouble with movement, which is often seen in advanced Parkinson’s disease during freezing episodes or when medication effects fluctuate.

**Fig. 14:**
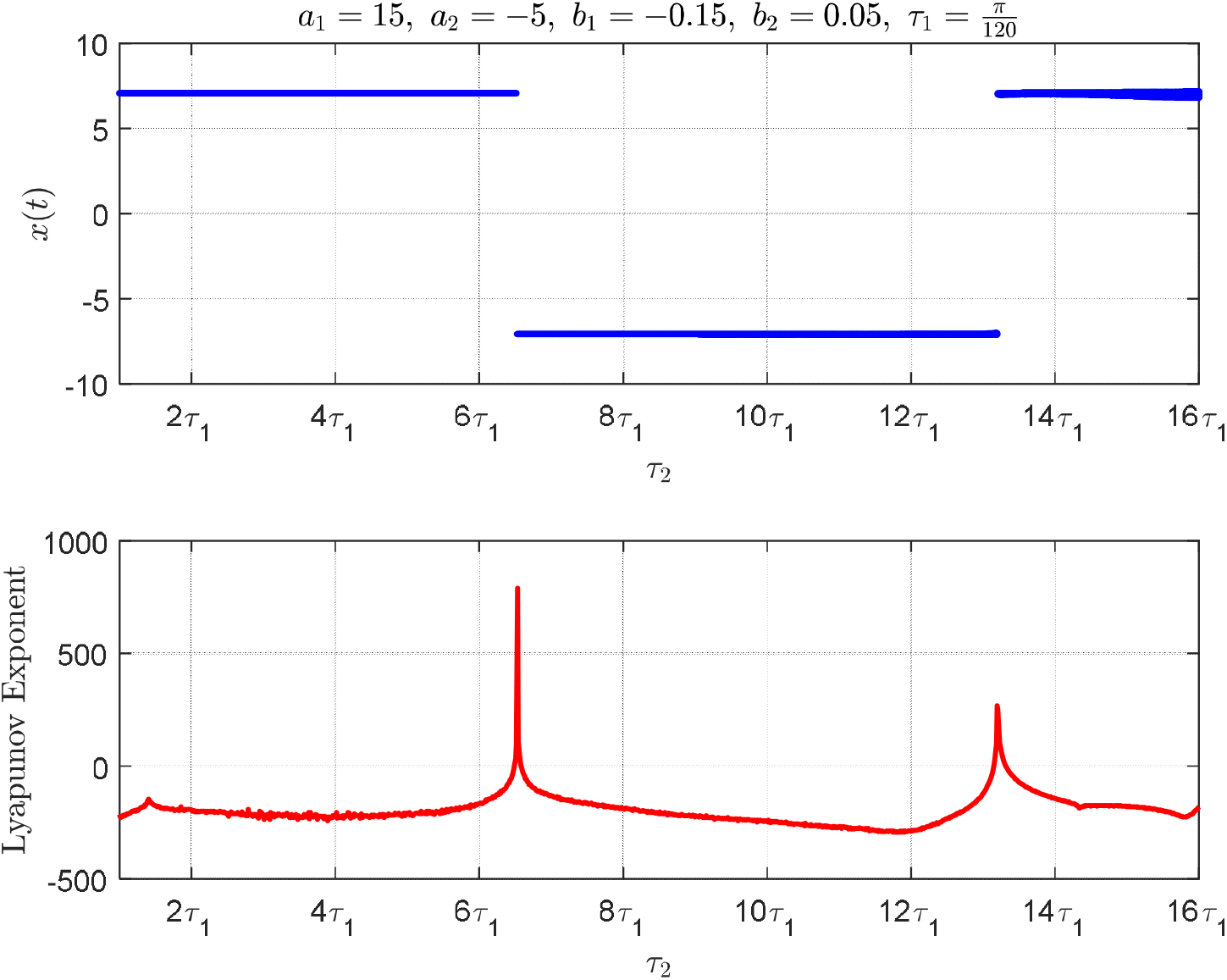
With negative saturation in both pathways, cause abrupt transitions between discrete stable states, Final 5% *of a* 5 − *second* simulation.

Figure 15 investigates the impact of the simulation time horizon using two durations: *t*_*final*_ = 0.5 *s* and *t*_*final*_ = 1.5 *s*, under the parameters *a*_1_= 15, *a*_2_ = − 5, *b* = − 0.30 and *b*_2_ = 0.10. Initially, the system exhibits either subthreshold activity or excessive excitation depending on the delay in the indirect pathway. With extended simulation, transient dynamics gradually stabilize, revealing convergence toward consistent trajectories. This behavior matches the motor symptoms seen in Parkinson’s disease, where slow movement or hyperactivity happens because of an imbalance in delayed signals that excite and inhibit the system (Fig. 15).

**Fig. 15:**
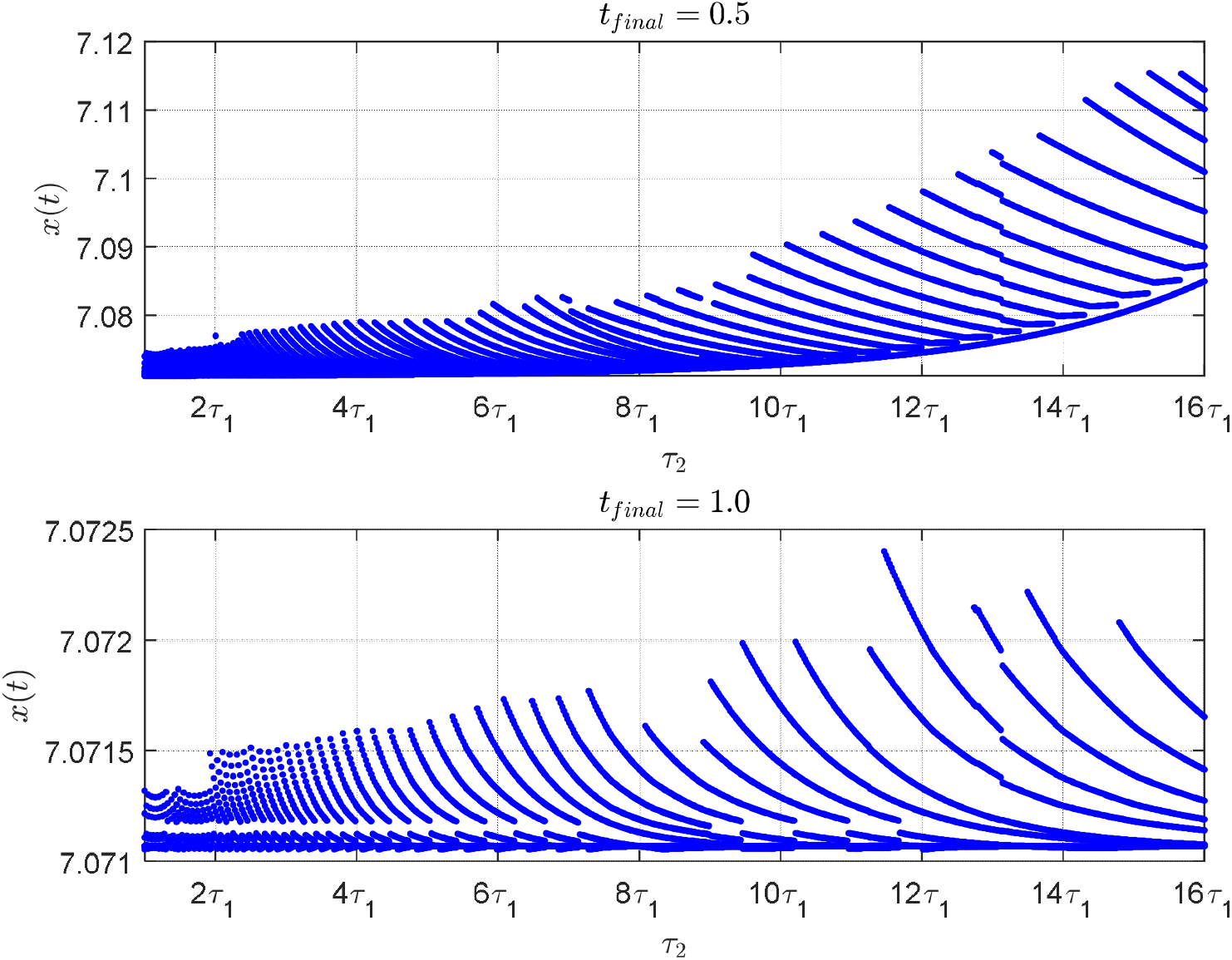
Time-dependent bifurcation patterns under moderate indirect excitation. At short durations (0.5 s and 1.0 s). Final 5% of each simulation.

Figures 7-15 illustrate how crucial the nonlinear saturation parameters _1_ and _2_, along with the length of the simulation, are in influencing the behavior of delayed neural feedback systems. The different outcomes—from being unstable to having controlled chaos and several stable conditions—demonstrate that the model can imitate various movement problems, like tremors, freezing, dyskinesia, and bradykinesia. These insights affirm the value of time-delayed nonlinear models in capturing the complexity of neuromotor regulation. Changes between stable and unstable states are closely connected to *τ*_2_, which shows how the long-loop cortico-basal ganglia work together. So, adjusting the nonlinear saturation in feedback circuits could be a useful way to identify motor disorders and a promising focus for drug or brain stimulation treatments.

Figure 16 shows how the system acts over time and frequency when it is stable *a*_1_ = 0.30, *a*_2_ = 0.30, *b*_2_ = 0.15) while changing the secondary *τ*_2_ to different values (2*τ*_1_,4.2 *τ*_1_, 9 *τ*_1_, 14 *τ*_1_), with 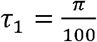. This setup shows a slightly curved increase in activity, which is good for simulating low-energy or resting states in cortico-basal ganglia circuits. In the top panel, the time-domain response x(*t*) shows that all four delay settings lead to rhythmic oscillations, but they have different heights and speeds of reaching those oscillations. For *τ*_1_ = 4.2*τ*_1_ (green), the system swiftly stabilizes into high-amplitude, low-frequency oscillations. On the other hand, when *τ*= 9*τ*_1_ and *τ*_2_ = 14*τ*_1_, the system shows smaller and more unpredictable changes, which relate to interference effects in longer feedback loops. Significantly, the *τ*_2_ =2*τ*_1_ scenario (blue) has a higher rapid damping envelope, signifying enhanced attenuation of early transients (Fig. 16).

**Fig. 16:**
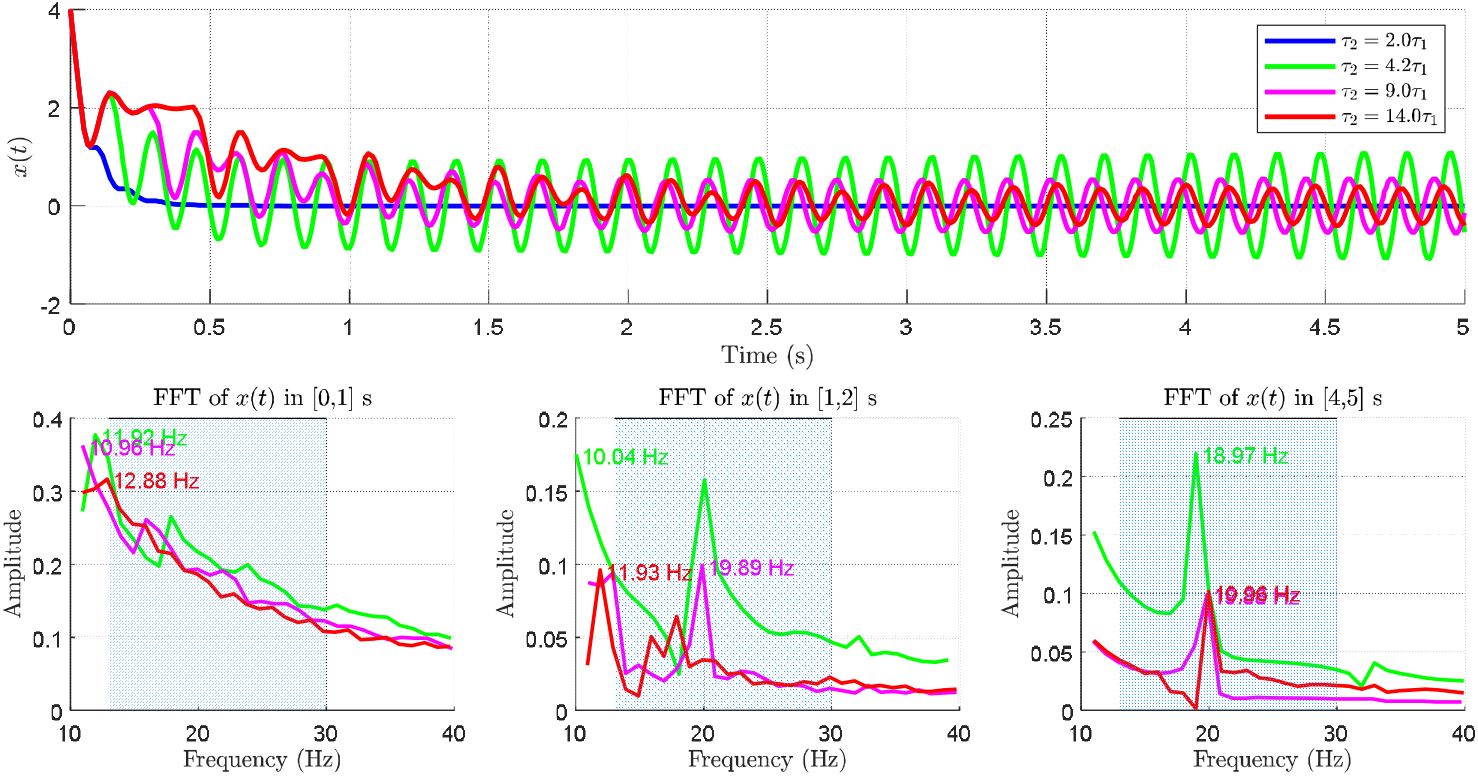
Oscillatory patterns evolve with indirect delay: time-domain traces show transient decay and stable rhythms, while frequency analysis across windows reveals delay-driven beta-band modulation.

The FFT analyses (bottom panels) illustrate the evolution of dominant frequencies throughout three temporal intervals: [0,1] *s*, [1,2] *s* [4,5) *s* and. In the initial phase ([0,1] *s*), all traces display broadband power within the 10 -– 25 *Hz* range, with peak frequencies at approximately 12.9 *Hz*(red), 11.9 *Hz* (green), and 13.6 *Hz* (magenta). These peaks indicate the system’s temporary convergence dynamics prior to the establishment of frequency locking.

During the second interval [1,2] *s*, the spectrum constricts, disclosing a higher number of coherent oscillatory states. The green trace (*τ*_2_ = 4.2*τ*_1_) stabilizes at approximately 19.93 *Hz*, whereas the other variants (magenta and red) have broader profiles with peaks beneath 13 *Hz*.This study shows that medium delay values *τ* = 4.2*τ* help the system strongly connect to the beta band, which is a feature of brain activity when at rest.

By the concluding interval [4,5], the system distinctly differentiates into two dynamic regimes. The green trace exhibits a consistent beta peak (∼19.9 H*z*), signifying resonance locking. The red and magenta curves exhibit diminished spectral power and lowered frequency peaks, indicating subharmonic or detuned responses. The blue trace (*τ*_2_ = 2*τ*_1_) is nearly entirely attenuated, exhibiting minimum spectral content.

The results confirm that the delay *τ*_2_ serves as a crucial bifurcation parameter, affecting both the amplitude and spectral properties of the system. Specifically, delays of about 4.2*τ*_2_ promote the development of steady low-beta oscillations, a pattern frequently linked to optimal rest-state basal ganglia functionality (Fig. 16). Longer delays because the system has chaotic or weaker oscillations, which is similar to the disorganized rhythms found in diseases like Parkinson’s. This analysis emphasizes the value of timing coordination in feedback circuits and supports the use of the amplitude and spectral properties of the system. Specifically, delays of about 4.2*τ*_2_ -based tuning in adaptive stimulation paradigms aimed at stabilizing motor control networks.

## 4. Results and Discussion

The dynamic behavior of the corticobasal ganglia loop was systematically analyzed. Each state shows how the network interacts based on the strengths and timing of the direct and indirect pathways. Collectively, the results shed light on how time delays in feedback loops contribute to the emerging beta-band synchronization, a hallmark of Parkinson’s disease.

The system dynamics in the resonance study (Figs. 1–3) show large fluctuations when the indirect pathway’s delay (the amplitude and spectral properties of the system. Specifically, delays of about (*τ*_2_) is small, but these fluctuations become more stable as _2_ gets larger. These oscillations are dampened by interactions between negative saturation in the direct pathway and positive saturation in the indirect pathway.

However, looking at resonance in this way did not provide enough information about the resting, active, and inhibited states of the motor cortex in the corticobasal ganglia loop. So, to explore the three functional states, we used bifurcation analysis with longer synchronization time between the two pathways (*τ*_2_) and Lyapunov analysis, as shown in Figures 4–15.

- In the resting state (Figures 4 to 6), the simulations show that as □ increases relative to *τ*;[, transitions to chaotic behaviors occur with fluctuations of varying intensity, which are consistent with the onset of resting tremor in Parkinson’s disease, due to impaired feedback timing.
- In the case of being overly excited or active (Figs. 7-11), the simulations showed that when *τ*_2_ is high, the motor cortex becomes overly active, then slows down (bradykinesia), and then becomes active again, which matches the advanced symptoms of Parkinson’s disease. If the activity becomes too high because of complex interactions, the direct pathway steps in to stop the motor cortex from activating right away, acting as a safety measure that only kicks in during times of hyperactivity.
- In the inhibited state (Figures 12–15), activity did not reach the threshold required to trigger movement, indicating a delayed motor response at the onset of movement, followed by excessive activity, a pattern seen clinically in Parkinsonian motor disorders.

Looking at Figures 4-6 and 7-15 shows that symptoms usually start with shorter resting delays, which supports the idea that tremors happen first, followed by slow movement and freezing as dopamine loss gets worse. Notably, the “freezing” pattern appeared at *τ*_2_ ≈ 6.5*τ*_1_ in inhibited states, reflecting the nature of symptom progression in advanced disease.

The results in figure 16 show that resonance happens in the beta frequency range (13 - 30 *Hz*–), highlighting how the timing of feedback through the indirect pathway is linked to the development of harmful oscillations.

We propose that the striatum, the first major input point into the basal ganglia, is the origin of the initial dysfunction of the basal-cortical loop. Timing abnormalities in this component generate resting-state beta oscillations. These oscillations then interact with nonlinear interactions between the subthalamic nucleus and the globus pallidus, exacerbating motor symptoms. In particular, the findings suggest that the globus pallidus may play a crucial role in reshaping the dynamic gain of the indirect pathway, thereby altering the transition from tremor to bradykinesia and freezing.

## 6. Conclusion

This study developed and analyzed a dual-delay nonlinear model of the cortico-basal ganglia circuitry, inspired by the van der Pol oscillator, to investigate the origins and dynamics of pathological motor rhythms in Parkinson’s disease. By incorporating separate feedback delays for the direct and indirect pathways, the model successfully captured key motor phenomena such as rest tremor, bradykinesia, dyskinesia, and freezing of gait.

Through systematic bifurcation and Lyapunov analyses, we demonstrated that the emergence and evolution of these symptoms are strongly dependent on the ratio and magnitude of time delays, as well as the nonlinear saturation effects in both pathways. Specifically, it was shown that the ratios between the time delays and the signal strength in each path lead to internal resonance. As delays increase and nonlinear interactions intensify, the system transitions through a spectrum of dynamic states, including stable rhythms, quasi-periodicity, and chaos— mirroring the clinical progression from mild tremor to severe motor inhibition.

A major insight from our findings is the origination of pathological dynamics in the striatal structures, with progressively worsening effects mediated by the globus pallidus, particularly under nonlinear conditions. This offers a mechanistic explanation for the sequence of symptoms observed in Parkinson’s disease, highlighting how disrupted excitation-inhibition timing in feedback circuits leads to delayed movement initiation or excessive suppression.

The model suggests that targeted modulation of time delays—either pharmacologically or through neural activity regulation techniques such as adaptive deep brain stimulation—could be used to disrupt pathological resonance and restore normal motor function. This model provides a coordinated, physiologically sound framework for understanding delay-induced motor disturbances in Parkinson’s disease. It links theoretical insights from nonlinear dynamics systems with real-world clinical phenomena, opening new avenues for improved therapeutic strategies based on tuning the timing of neurofeedback.

## Authors Statements

**M. A. Elfouly:** Investigation, Resources, Methodology, Validation, Conceptualization, Data duration, Data duration, Reviewing and Editing.

**T. S. Amer:** Formal analysis, Validation, Visualization and Reviewing.

## Funding Acknowledgement

No specific funding was provided for this research by any public, commercial, or not-for-profit organization.

## Data Availability

All data generated or analyzed in this study are included in this published article.

## Ethical Approval

There is no clinical data. human or animal experiments. The manuscript’s data were suggested by the author to explain the model’s behavior and the generation of neural impulses.

## Availability of data and materials

The datasets used and/or analysed during the current study available from the corresponding author on reasonable request.

